# Megakaryocytes, erythropoietic and granulopoietic cells express CAL2 antibody in myeloproliferative neoplasms carrying CALR gene mutations

**DOI:** 10.1101/2019.12.22.886317

**Authors:** Hebah Ali, Ignazio Puccio, Ayse U Akarca, Roshanak Bob, Sabine Pomplun, Wai Keong Wong, Rajeev Gupta, Mallika Sekhar, Jonathan Lambert, Hytham Al-Masri, Harald Stein, Teresa Marafioti

**Affiliations:** Haematological Malignancy Diagnostic Service, Leeds; University of Leeds, Leeds; Department of Histopathology, University College London, London; Reference and Consultation Center for Lymphoma and Haematopathology, Pathodiagnostik Berlin, Berlin; Department of Cellular Pathology, University College Hospital, London; Department of Haematology, University College Hospital, London; Hematogenix Laboratory Services, Chicago

**Author notes:** Equal contribution (Ignazio Puccio, Ayse U Akarca). Equal senior authorship (Harald Stein, Teresa Marafioti).

**Keywords:** mutated Calreticulin, CALR, CAL2, myeloproliferative neoplasms, myelofibrosis, essential thrombocythemia

## Abstract

The discovery of mutated Calreticulin (CALR) in myeloproliferative neoplasms (MPN) has provided proof of clonality, diagnostic importance, and influence on prognosis of this pathology. The identification of this MPN-associated driver mutation -currently based on molecular assays- is represented as a major diagnostic criterion for essential thrombocythaemia (ET), pre-fibrotic myelofibrosis and primary myelofibrosis (PMF) in the updated World Health Organization (WHO) 2008 classification. In the present study, we validated by immunohistochemistry the diagnostic usefulness of the monoclonal CAL2 antibody. Cases of acute myeloid leukaemia (AML) and myelodysplastic/ myeloproliferative neoplasms (MDS/MPN) have been also investigated to assess the specificity of CAL2 antibody. For this purpose, the result of the CAL2 immunostaining was compared with the result of molecular assays. Additionally, we investigated by double staining whether expression of mutated CALR can also be demonstrated on cells of the erythroid and myeloid lineage. We confirmed the usefulness of the CAL2 monoclonal antibody in successfully detecting mutant CALR in bone marrow biopsies. We showed that the immune-reactivity of CAL2 was absolutely restricted to the presence of CALR mutations, which were seen only in ET and MDS/MPN biopsies, but not in AML biopsies (14/14). There was 100% concordance in biopsy specimens with the concomitant molecular results. We applied double staining technique and confirmed that a subpopulation of granulopoietic and erythropoietic cells express mutated CALR as demonstrated with the CAL2 antibody in cases of MPNs. This supports the suggestion that the CALR mutations occur in a multipotent progenitor capable of generating both myeloid and erythroid progeny with preferential expansion of megakaryocytic cell lineage as a result of CALR mutation in an immature hematopoietic stem cell.

## Introduction

The discovery of mutated Calreticulin (CALR) in myeloproliferative neoplasms (MPN) has provided proof of clonality, diagnostic importance, and influence on prognosis of this pathology. The identification of this MPN-associated driver mutation -currently based on molecular assays- is represented as a major diagnostic criterion for essential thrombocythaemia (ET), pre-fibrotic myelofibrosis and primary myelofibrosis (PMF) in the updated World Health Organization (WHO) 2008 classification ^[1]^. Recently, Vannucchi et al ^[2]^ and Stein et al ^[3]^ raided polyclonal and monoclonal antibodies against mutated CALR to be used in the routinely processed bone marrow paraffin sections as a complimentary assay to detect the mutant CALR protein in patients with MPNs. In the present study, we validated by immunohistochemistry the diagnostic usefulness of the monoclonal CAL2 antibody. Cases of acute myeloid leukaemia (AML) and myelodysplastic/ myeloproliferative neoplasms (MDS/MPN) have been also investigated to assess the specificity of CAL2 antibody. For this purpose, the result of the CAL2 immunostaining was compared with the result of molecular assays. Additionally, we investigated by double staining whether expression of mutated CALR can also be demonstrated on cells of the erythroid and myeloid lineage.

## Material and Methods

### Tissue Samples

One hundred and eighty two bone marrow biopsies from patients with myeloid neoplasms: series of MPNs (n=66) and a control group of acute myeloid leukaemia (AML) (n=116) were retrieved from the files of the Department of Histopathology at University College London Hospitals. The cases were diagnosed by expert haematopathologists and haematologists at University College London Hospitals following the criteria of the 2008 WHO classification ^[1]^. Approval for this study was obtained from the National Research Ethics Service, Research Ethics Committee 4 (REC Reference number 09/H0715/64).

### Immunohistochemistry

The bone marrow biopsies were fixed in 10% neutral buffered formalin, decalcified for six hours using Gooding Stewart solution, processed and embedded in paraffin. Immunostaining using the newly developed anti-human CAL2 monoclonal mouse antibody ^[3]^ was performed on bone marrow biopsies tissue sections using the Roche-Ventana BenchMark ULTRA autostainer (Ventana Medical Systems, Tucson, US).

The CAL2 antibody was assessed under different conditions (i.e. dilution and antigen retrieval protocols) and the chosen dilution, which showed selective background-free reaction was 1:10. Counterstaining was performed using haematoxylin and bluing reagent from Ventana/Roche. Slides were mounted with cover slips and air-dried.

### Double Immunostaining

Double immuno-enzymatic labelling of biopsies’ sections following the described pre-treatment was carried out by means of the EnVision peroxidase and alkaline phosphatase kits (DakoCytomation). Primary antibodies were incubated for 30 minutes at room temperature, and a diaminobenzidine (DAB) substrate (DakoCytomation) was then used for detection of antibody binding. Sections were then incubated for 30 minutes with the second antibody, and then for an additional 30 minutes with the alkaline phosphatase EnVision kit (DakoCytomation). The second reaction was detected by means of a Vector blue alkaline phosphatase substrate kit (Vector Laboratories, Peterborough, UK). The sections were washed in tap water and mounted in aquamount (Merck, Poole, UK). Double immunostaining was carried out to analyse the expression of CD71, myeloproxidase, CD61 and GATA-1 in combination with CAL2 antibody. The cases were reviewed by an expert haematopathologist (TM), a co-author of this paper.

### Molecular Assay

For some cases included in this study, molecular analysis of CARL gene and a set of other genes including JAK2, MPL, BCR/ABL1 and KIT were performed using conventional DNA PCR. DNA was isolated from peripheral blood or bone marrow aspirate. DNA was amplified using forward and reverse primers spanning exon 9 of the CALR gene with the forward primer fluorescently labelled. CALR F: (5’ FAM-GGCAAGGCCCTGAGGTGT 3’) and CALR R: (5’ - GCCTCAGTCCAGCCCTG 3’). The conditions were: (1×) 95.0°C 15 min. (10×) 94.0°C 15 sec, 55.0°C 15 sec, 72.0°C 30 sec. (20×) 89.0°C 15 sec, 55.0°C 15 sec, 72.0°C 30 sec. (1×) 72.0°C 10 min. PCR products were analysed by capillary gel electrophoresis. 100 ng of gDNA extracted using QIAamp DNA Blood Mini Kit (Qiagen) was used in each assay.

## Results

CAL2 immunostaining was evaluated in a total of one hundred and eighty two bone marrow biopsies from patients with myeloid neoplasms (AML n=116, MPNs n=66) [Table 1]. Positivity was observed in twenty out of sixty-six biopsies from patients with MPNs that included: ET n=14, chronic MPN, with MF n=3; MDS/MPN, unclassifiable n=2; MF n=1 [Table 2]. In 14 of the 20 CAL2 positive cases, PCR or sequencing was performed and results showed CALR molecular aberrations (either mutation or deletion etc.) [Table 2].

Mutated CALR expressions shown with CAL2 was mostly restricted to megakaryocytes, principally labelling the cytoplasm and displaying a granular staining pattern [Figure 1]. Even some small CAL2 positive cells proved to be positive for CD61 thus identifying these cells as small megakaryocytes [Figure 2]. However, occasional smaller cells with round nuclei were stained by CAL2 antibody and antibodies to myeloperoxidase or to CD71, showing that a few granulopoietic (myeloproxidase positive) and erythropoietic (CD71 positive) cells express mutated CARL2. None of the acute myeloid leukaemia biopsies showed a positive staining with the CAL2. Double staining showed the CAL2 positive megakaryocytes in essential thrombocythaemia patient to co-express GATA-1 [Figure 1].

**Figure 1 caption:**
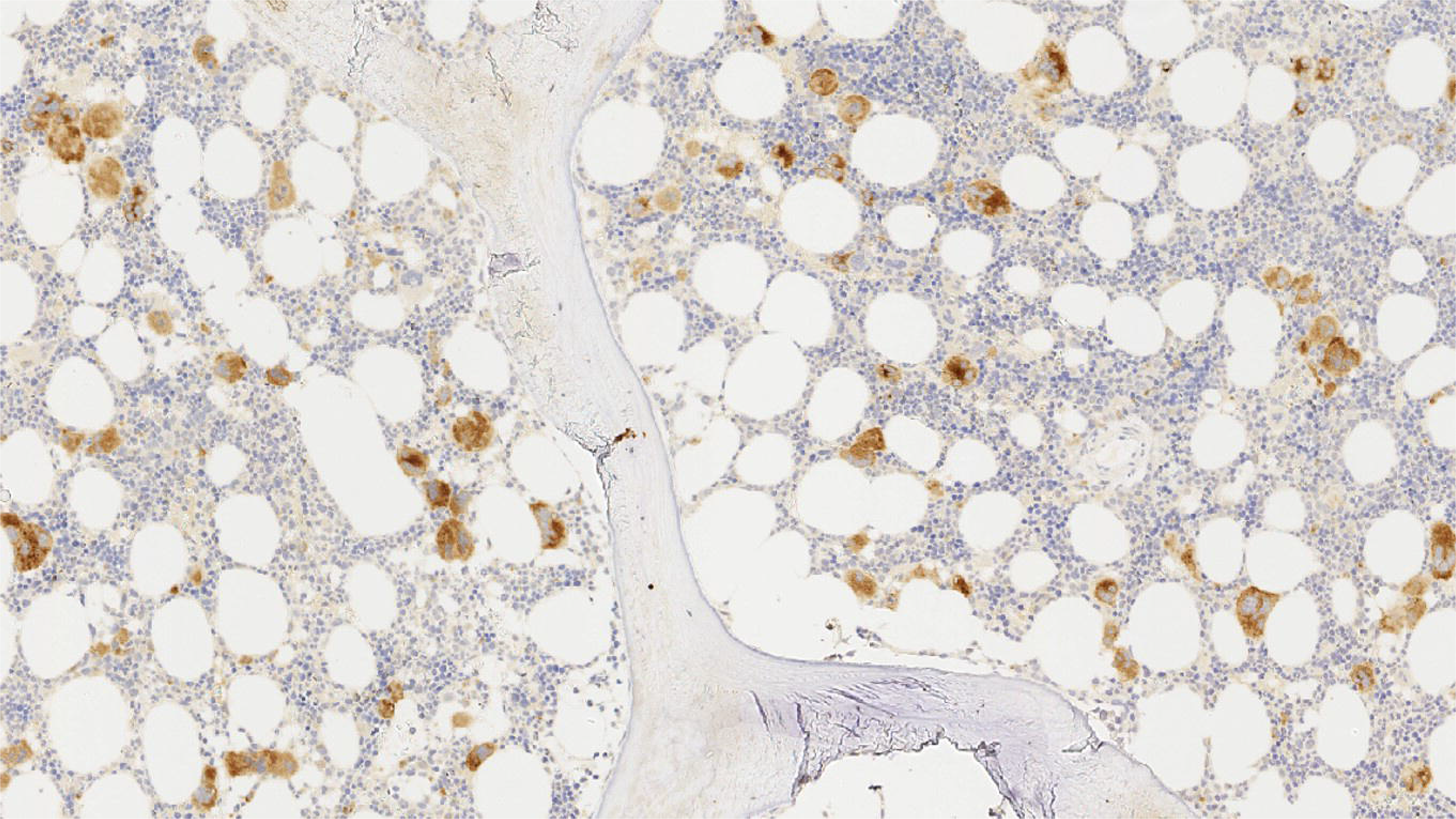
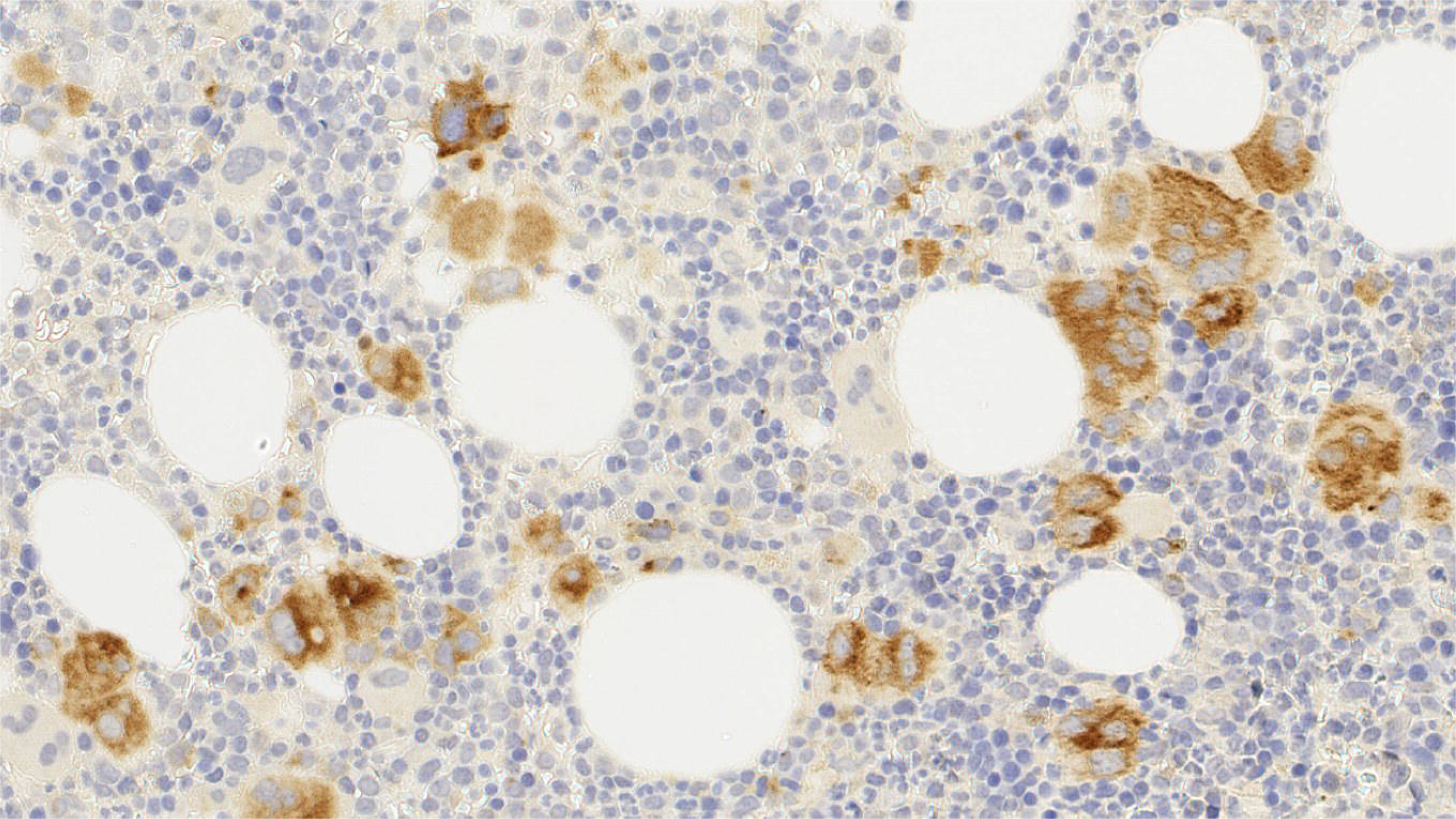
Imunostaining with CAL2: The megakaryocytes show strong expression in bone marrow biopsies from (a,b) an essential rombocythaemia and (c,d) a primary myelofibrosis patients. Magnification: ×4, ×20.

**Figure 2 caption:**
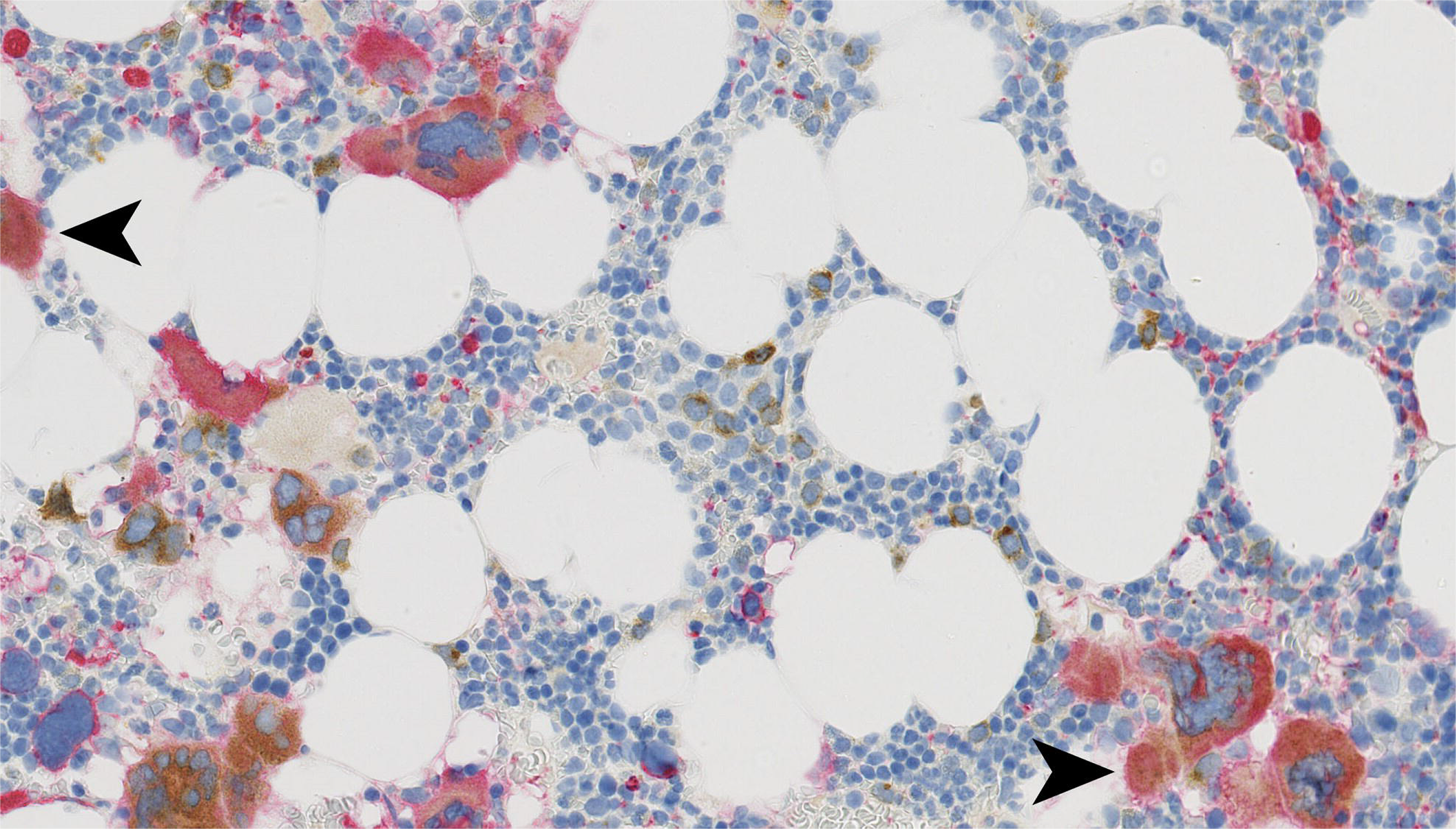
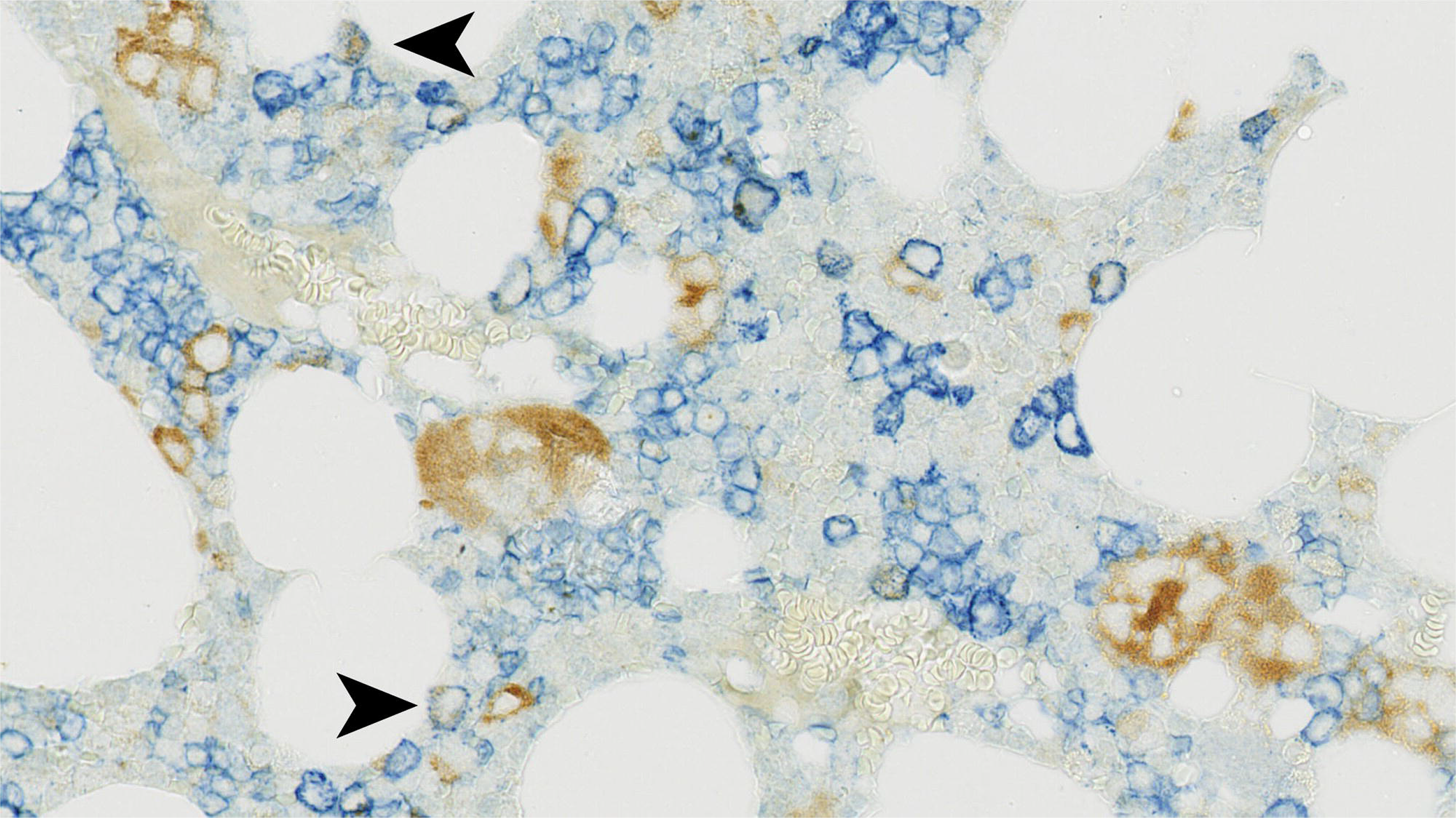
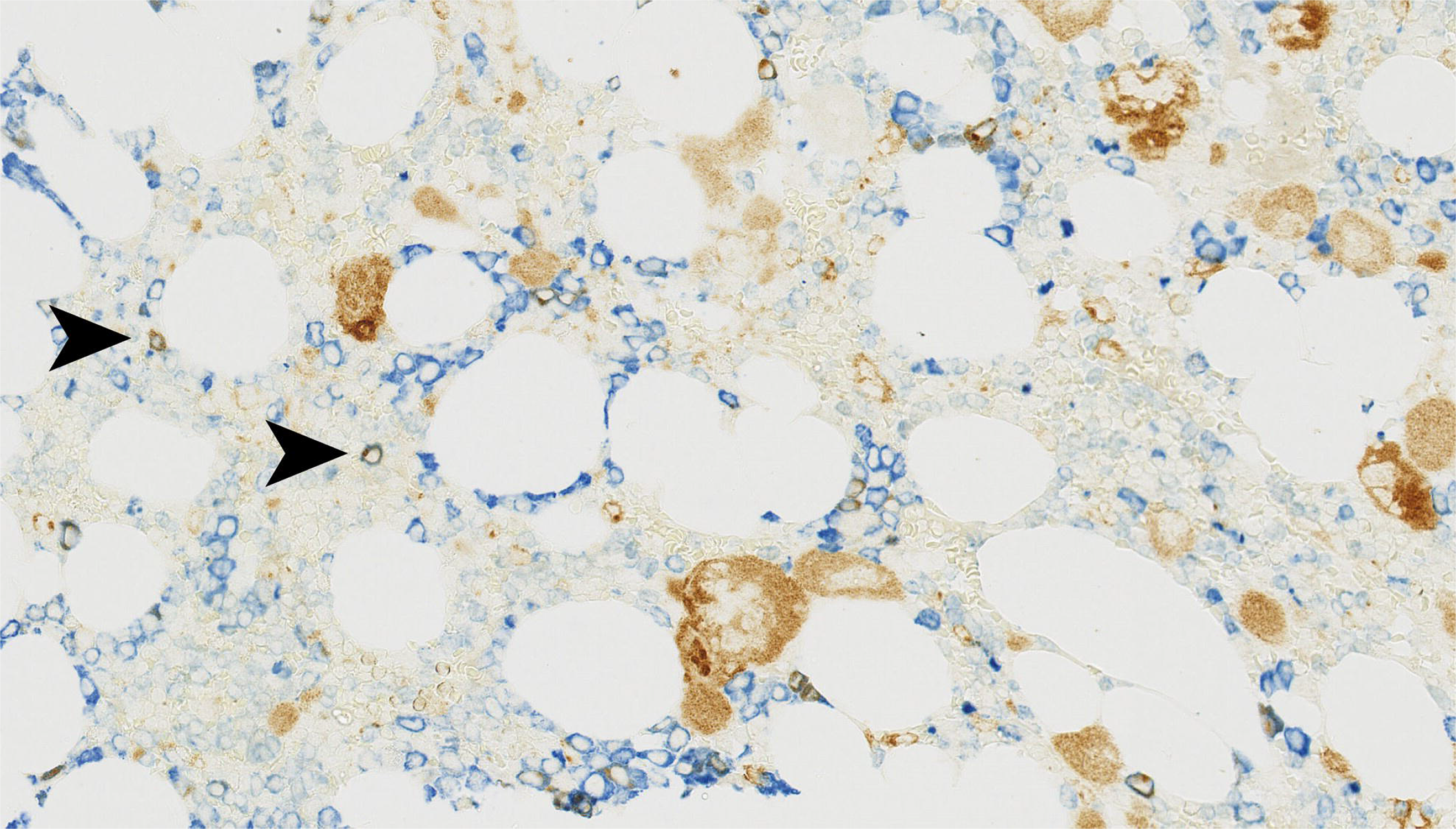
Double immunostaining with CAL2 and CD61, CD71 and MPO: CAL2 (brown cytoplasmic) is expressed in megakaryocytes well as smaller cells: (a) micro-megakaryocytes:CD61 red cytoplasmic, (b) erythroid precursors: CD71 blue cytoplasmic d (c) myeloid elements: myeloperoxidase blue cytoplasmic. Magnification: ×40.

**Figure 3 caption:**
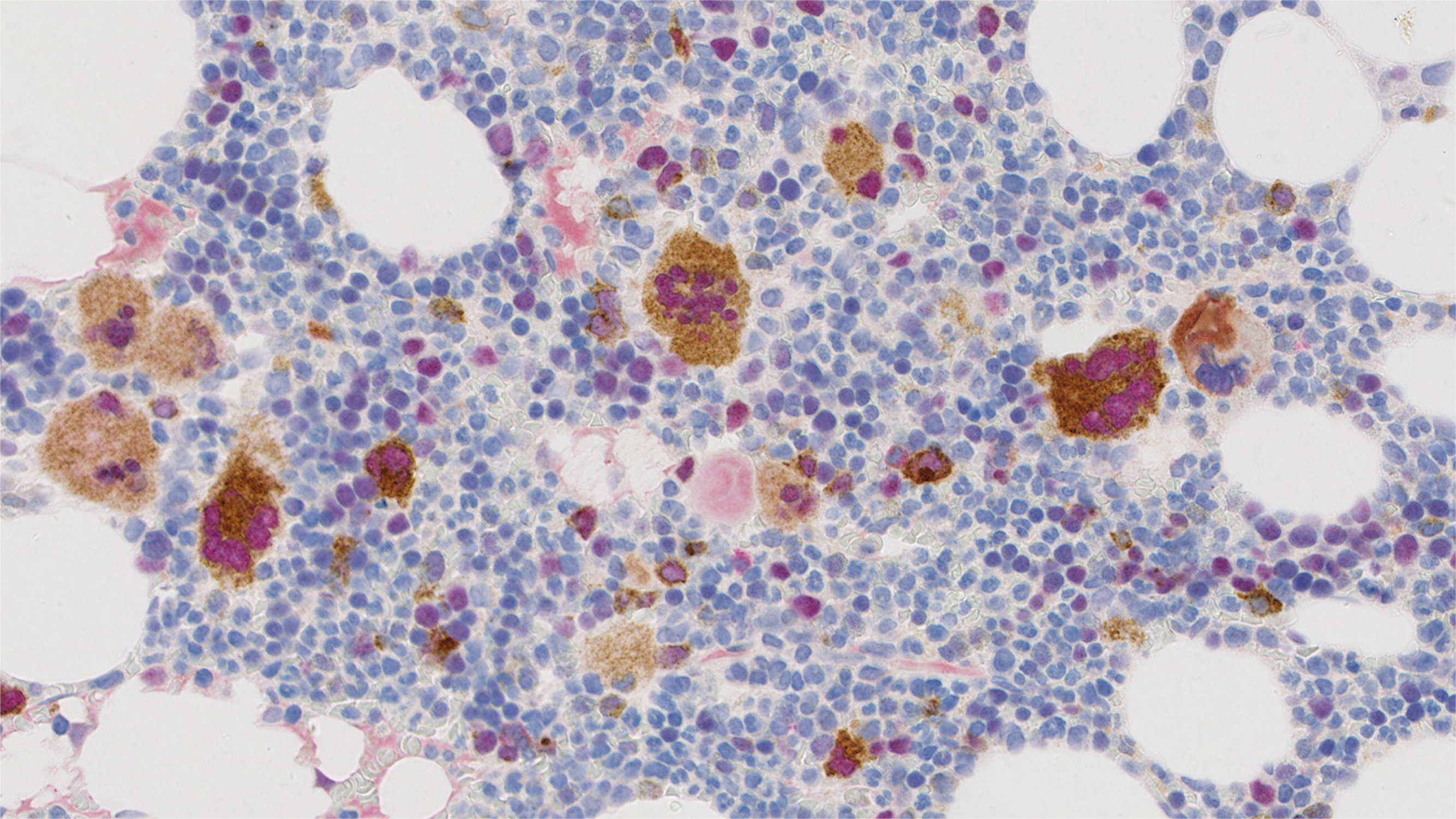
Double immunostaining with CAL2 and GATA-1: The megakaryocytes show double expression of CAL2 (brown cytoplasmic) d GATA-1 (red nuclear). Magnification: ×40).

## Discussion

Mutations in CALR have been discovered in 50% to 80% of JAK2 and MPL wild-type patients with Philadelphia-negative MPNs. The recent revision of WHO criteria for ET and PMF includes testing for the CALR mutation ^[1]^. Currently, molecular methods mainly by PCR are the standard for detection of these mutations. Unlike JAK2 and MPL point mutations, CALR mutations are highly heterogeneous, with several types of insertions or deletions, all located in exon 9 ^[4]^. Because of this high heterogeneity, the molecular assays are complicated. In addition, their performance might be time consuming, and technically or financially not feasible to many routine diagnostics labs. There is therefore a need for a simpler, faster cost-effective method. Recently, two groups worked on developing both polyclonal and monoclonal immunohistochemical stains to detect mutant CALR in bone marrow biopsies ^[2,3]^.

In our study, we confirmed the usefulness of the CAL2 monoclonal antibody in successfully detecting mutant CALR in bone marrow biopsies.

We showed that the immune-reactivity of CAL2 was absolutely restricted to the presence of CALR mutations, which were seen only in ET and MDS/MPN biopsies, but not in AML biopsies. There was 100% concordance in biopsy specimens with the concomitant molecular results. Our findings support the results of a previous analysis performed by Stein et al demonstrating the diagnostic utility of the CAL2 antibody ^[3]^ and are comparable to four following studies. ^[5–8]^

Moreover, we were able to explore the lineage of few smaller cells expressing mutated CALR described in some of those studies. Interestingly, Vannucchi et al ^[2]^ observed modest labelling by a polyclonal antibody in myeloid and erythroid cells. However, the lineage of similar cells could not be clarified in the first study assessing the monoclonal CAL2 antibody ^[3]^.The results from subsequent publications were diverse. In one 2016 study, the nine positive cases show staining in the majority of megakaryocytes with little or no staining in any other cell types ^[5]^.On the other hand, Nomani et al. in the same year noted staining of small mononuclear in CALR mutant cases ^[6]^. By performing double immunofluorescence staining, they proposed that the small cells appeared to be myeloid cells or blasts. Recently, Bonifacio et al ^[7]^ published unifying results describing two different patterns of CAL2-positive staining. Pattern A is characterised by staining of almost only megakaryocytes. In contrast, there is staining of megakaryocytes and small elements at least partially being myeloid precursors in pattern B.

In our study, we applied double staining technique and confirmed that a subpopulation of granulopoietic and erythropoietic cells express mutated CALR as demonstrated with the CAL2 antibody in cases of MPNs. This supports the suggestion by Nanglia et al ^[9]^ that the CALR mutations occur in a multipotent progenitor capable of generating both myeloid and erythroid progeny with preferential expansion of megakaryocytic cell lineage as a result of CALR mutation in an immature hematopoietic stem cell.

Additionally using double staining, we detected co-expression of GATA-1 in ET biopsies. This is consistent with the finding of Rinaldi et al ^[10]^ and has relevance for the understanding of pathogenetic mechanisms associated with CALR mutations. Although at present, no direct correlation between GATA1 expression and CALR mutations is found, Brown et al observed a significant up regulation of CALR mRNA in MPN cases with high GATA-1^[11]^ and this could be relevant to the double immunohistochemistry staining in our study. This correlation should be confirmed in wider and independent series.

In conclusion, immunohistochemistry is readily available in the majority of diagnostic laboratories and the detection of CALR mutations by the CAL2 antibody represents a valuable supplement to traditional mutation testing and can help to facilitate the timely, appropriate selection and treatment of patients with myeloproliferative neoplasms with targeted therapies.

## Conflict of Interest

The authors have no conflict of interests.

## Acknowledgments

This study was funded and supported by the National Institute for Health Research University College London Hospitals Biomedical Research Centre (TM) and Cancer ImmunoTherapy Accelerator (CITA) CRUK

## Authorship

TM and HS conceived the idea and designed the study. TM, MS and JL selected the clinical cases for inclusion. PI and AUA performed immunostaining and collate the data. TM reviewed and interpreted the staining. TM and HA compiled the results and created the figures. HM, WKW, RG, MS, and JL provided a substantial number of clinical cases. WKW, RG and SP involved in the diagnostic analysis of the cases. HA and TM wrote the manuscript.

