## Supplementary material for "Megakaryocytes, erythropoietic and granulopoietic cells express CAL2 antibody in myeloproliferative neoplasms carrying CALR gene mutations": Tables 1 and 2

Table 1

| Diagnosis | Total number | CAL2 staining (positive/total) |
|---|---|---|
| AML | 116 | 0/116 |
| MPN | 66 | 20*/66 |

Table 1 legend:

CAL2 immunohistochemistry staining results in bone marrow biopsies obtained from patients with myeloproliferative neoplasms and acute myeloid leukaemia

Abbreviations: AML: acute myeloid leukaemia. MPN: myeloproliferative neoplasms

* The twenty CAL2 positive cases are detailed in Table 2.

Table 2

| ID | P/R | Diagnosis | Molecular CALR aberrations/ others | CAL2 IHC |
|---|---|---|---|---|
| ID1 | P | ET | 2 bp insertion in exon 9 and a c.149_1154del and ins TCCTTGTC resulting in predicted protein change p. (Glu383Aspfs*48). | + |
| ID2 | P (2012) | ET | Not performed | + |
| ID2.1 | R (2014) | Chronic MPN with MF | 5 bp insertion in exon 9. The mutation load was approx. 47%. The mutation c.1154_1155ins TTGTC resulting in predicted protein change p. (Lys385Asnfs*47). | + |
| ID3 | P | ET | Not performed | + |
| ID4 | P | ET | Not performed | + |
| ID5 | P | ET | Not performed | + |
| ID6 | P | MDS/ MPN, unclassifiable | CALR exon 9 mutation | + |
| ID7 | P | ET | Not done | + |
| ID8 | P | MDS/ MPN, unclassifiable | 52 bp deletion, the mutation load was 45%. The mutation c.1099_1150del resulting in predicted protein change p. (Leu367Thrfs*46). | + |
| ID9 | P | ET | Not performed | + |
| ID10 | P | ET | 52 bp deletion, the mutation load was 32%. The mutation c.1099_1150del resulting in predicted protein change p. (Leu367Thrfs*46). | + |
| ID11 | P | Chronic MPN with MF | Not performed | + |
| ID12 | P | ET / early stage MF | CALR+, BCR/ABL (-), JAK2 (-), MPL (-), KIT (-) | + |
| ID13 | P | Chronic MPN with MF | CALR+, BCR/ABL (-), JAK2 (-), MPL (-), KIT (-) | + |
| ID14 | P | ET / early stage MF | CALR+, BCR/ABL (-), JAK2 (-), MPL (-), KIT (-) | + |
| ID15 | P | ET / early stage MF | CALR+, BCR/ABL (-), JAK2 (-), MPL (-), KIT (-) | + |
| ID16 | P | ET | CALR+, BCR/ABL (-), JAK2 (-), MPL (-), KIT (-) | + |
| ID17 | P | ET | CALR+, BCR/ABL (-), JAK2 (-), MPL (-), KIT (-) | + |
| ID18 | P | Early stage MF | CALR+, BCR/ABL (-), JAK2 (-), MPL (-), KIT (-) | + |
| ID19 | P | MF | CALR+, BCR/ABL (-), JAK2 (-), MPL (-), KIT (-) | + |
| ID20 | P | ET | CALR+, BCR/ABL (-), JAK2 (-), MPL (-), KIT (-) | + |

Table 2 legend

Clinical, pathological and molecular data of myeloproliferative cases with CAL2 positive immunostaining.

Abbreviations: P: primary diagnosis. R: relapse. ET: essential thrombocythemia. MPN: myeloproliferative neoplasm. MF: myelofibrosis. MDS/ MPN: myelodysplastic/ /myeloproliferative neoplasms.
